# The Biological Evaluation of Fusidic Acid and Its Hydrogenation Derivative as Antimicrobial and Anti-inflammatory Agents

**DOI:** 10.1101/311951

**Authors:** Pan-Pan Wu, Hao He, W. David Hong, Tong-Rong Wu, Su-Qing Zhao, Xi-Ping Cui, Ying-Ying Zhong, Dong-Li Li, Xue-Tao Xu, Zhao-Jun Sheng, Bo-Rong Tu, Min Gao, Jun Zhou, Stephen A. Ward, Paul M. O’Neill, Kun Zhang

## Abstract

Fusidic acid (WU-FA-00) is the only commercially available antimicrobial from the fusidane family that has a narrow spectrum of activity against Gram-positive bacteria. Herein, the hydrogenation derivative (WU-FA-01) of fusidic acid was prepared, and both compounds were examined against a panel of six bacterial strains. In addition, their anti-inflammation properties were evaluated using a 12-*O*-tetradecanoylphorbol-13-acetate (TPA)-induced mouse ear edema model. The results of the antimicrobial assay revealed that both WU-FA-00 and WU-FA-01 displayed a high level of antimicrobial activity against Gram-positive strains. Moreover, killing kinetic studies were performed, and the results were in accordance with the MIC and MBC results. We also demonstrated that the topical application of WU-FA-00 and WU-FA-01 effectively decreased TPA-induced ear edema in a dose-dependent manner. This inhibitory effect was associated with the inhibition of TPA-induced up-regulation of pro-inflammation cytokines IL-1β, TNF-α and COX-2. WU-FA-01 significantly suppressed the expression levels of p65, IκB-α, and p-IκB-α in the TPA-induced mouse ear model. Overall, our results showed that WU-FA-00 and WU-FA-01 not only had effective antimicrobial activities *in vitro*, especially to the Gram-positive bacteria, but also possessed strong anti-inflammatory effects *in vivo*. These results provide a scientific basis for developing fusidic acid derivatives as antimicrobial and anti-inflammatory agents.

## Introduction

Over the past few decades, the appreciation of the key role of inflammation in disease diagnosis, prevention and treatment has burgeoned (1, 2). Inflammation has been defined as a complex biological response of vascular tissues to different types of harmful stimuli (3, 4), such as damaged cells, irritants or pathogens. Inflammation has also been linked to the release of pro-inflammatory cytokines (5, 6), including tumour necrosis factor-alpha (TNF-α), interleukin-1β (IL-1β), interleukin-6 (IL-6) and cyclooxygenase-2 (COX-2), all of which could be a sign of many diseases (4, 7). Therefore, inflammation is a biological response wherein the organism attempts to remove the injurious stimuli and initiate the healing process for the tissue; thus, it could be regarded as a protective effect (4).

Currently, steroids and non-steroidal anti-inflammatory drugs are proverbially used in clinical application as effective therapeutic anti-inflammatory agents (4). Despite the widespread use of anti-inflammatory drugs, there may be some residual risks of inflammation and the side effects of their long-term oral administration (8), especially in infectious diseases, in which patients suffer from not only the inflammatory responses but also pathogenic microorganism infections (9–11).

Fusidic acid (FA, WU-FA-00) (Figure 1), which has a steroid-like scaffold structurally and is derived from the fungus *Fusidium coccineum*, is the only marketed antibiotic from the fusidane family. Sodium fusidate, the sodium salt of fusidic acid, was primary introduced into practice as an anti-staphylococcal therapy in 1962 (12–14). However, FA has a narrow spectrum of biological activity against some anaerobic gram-negative organisms and most gram-positive bacteria, especially the staphylococci, including the methicillin-resistant *Staphylococcus aureus* (MRSA) and coagulase-negative staphylococci (15–17). Although some antimicrobial activity and reasonable anti-inflammatory effects have been discovered (18, 19), there is no in-depth study of FA and its derivatives as potential anti-inflammatory agents. Therefore, the therapeutic efficacy of FA and its derivatives as antimicrobial and anti-inflammatory agents should be explored.

**Fig. 1.**
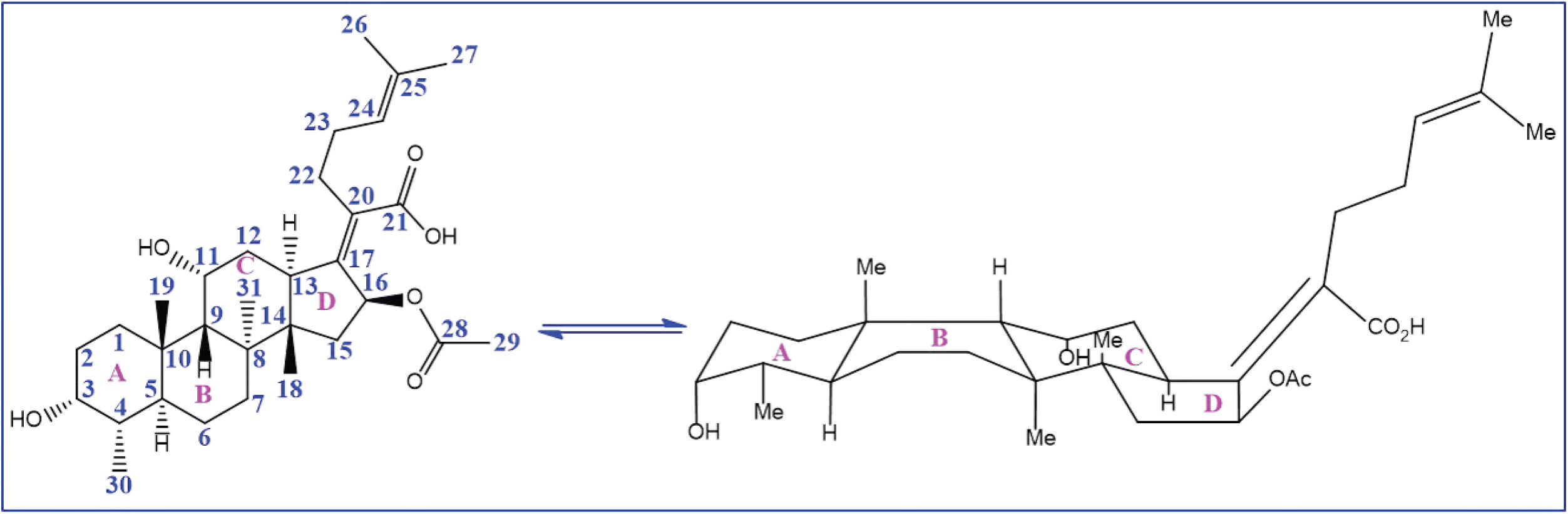
Chemical structure and conformation of fusidic acid (WU-FA-00).

In the present study, the *in vitro* antimicrobial activities of fusidic acid and its hydrogenation derivative (WU-FA-01) were investigated against six bacterial strains, including five Gram-positive bacteria strains and one Gram-negative bacteria strain. In parallel, 12-*O*-tetradecanoylphorbol-13-acetate (TPA) was used as an inducer to explore both compounds, inhibitory activity on skin inflammation in a mouse ear edema model (20–22). Moreover, immunohistochemical analysis was introduced to reveal their inhibitory effects on the expression of TPA-induced TNF-α, IL-1β and COX-2 in mouse ears. Furthermore, the anti-inflammatory mechanisms of FA and its hydrogenation derivative were also discussed to gain insight into their effects. Accordingly, fusidic acid or its derivatives, especially the hydrogenation compound, may be developed as promising di-functional parent drugs, which could be underlying anti-inflammatory and antimicrobial agents.

## Results

### Chemistry

To obtain the hydrogenation derivative of FA, structural modifications (according to a previous study) were made at the double bond position of C-24 and C-25 (23). The synthetic route is shown in Scheme 1. The 24, 25-dihydrofusidic acid (WU-FA-01) was prepared by Palladium catalysed hydrogenation in quantitative yielding. Its structure was confirmed by high-resolution mass spectrometry (HRMS), CHNS-O elemental analyser, ^1^H NMR and ^13^C NMR, and it was in accordance with the previous research (23).

### Antibacterial activity

The antibacterial activity of WU-FA-00 and WU-FA-01 were tested against six microorganisms, including reference strains consisting of Gram-negative bacteria and Gram-positive bacteria. All bacterial strains were cultured in Muller Hinton agar at 37 °C overnight.

### Agar disk diffusion method

The results of the antimicrobial activity of WU-FA-00 and WU-FA-01 against six different microorganisms are summarized in Table 1. Two different concentrations were examined in this method. The sizes of the inhibition zone indicated that the tested compounds with Gram-positive bacterial strains were larger than those with Gram-negative strains, and both compounds showed dose dependence. The inhibition zone diameter was in the range of 10.37±1.23 to 24.22±1.66 mm for Gram-positive strains. However, both WU-FA-00 and WU-FA-01 showed no inhibitory effect against the Gram-negative strains. Furthermore, the screening of the antimicrobial potential of the two compounds revealed that reducing the double bond to a single bond at positions C-24 and C-25 could retain their antimicrobial activities, specifically against the Gram-positive strains.

**Table 1.**
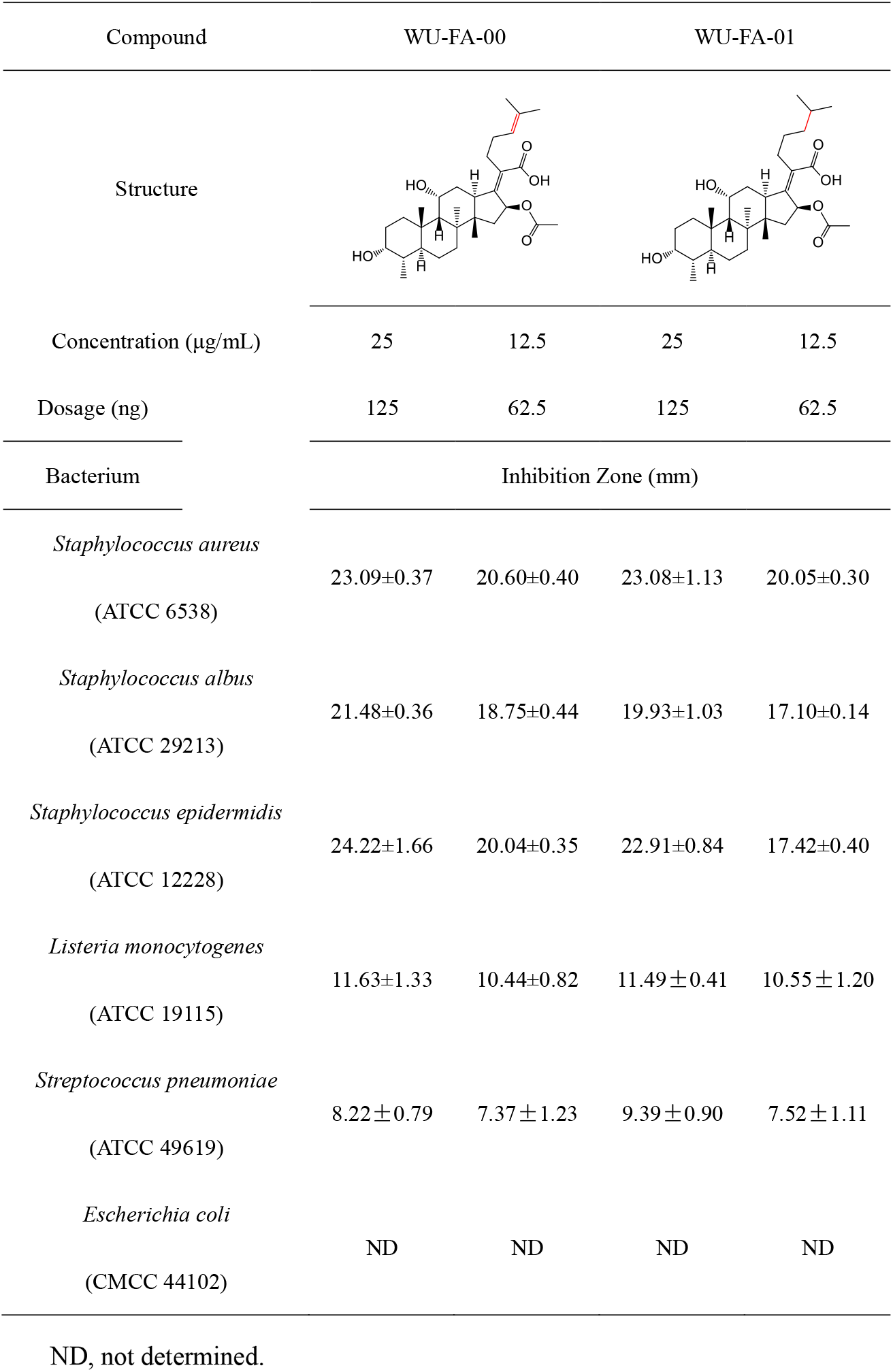
Antibacterial activity of WU-FA-00 and WU-FA-01 expressed in the inhibition zone (mm).

### Broth microdilution method

A microtiter plate dilution method was conducted to determine the minimal inhibitory concentration (MIC) and the minimal bactericidal concentration (MBC) in a 96-well plate. At the end of the incubation period, the plates were evaluated for the presence or absence of bacterial growth. Each sample concentration was tested four times against each microorganism. WU-FA-00, the parent compound, was employed as a positive control against bacterial growth. The final concentration of DMSO in the 96-plate well had no effect on bacterial growth.

WU-FA-00 and WU-FA-01, the two tested compounds, were found to be active against the microorganisms studied, especially the Gram-positive bacteria. The MIC and MBC values of the two compounds were determined according to the results of the micro-dilution method (Table 2). The results suggested that WU-FA-01 (MIC=100-625 ng/mL, MBC=200-1250 ng/mL) showed activity similar to its parent compound WU-FA-00 (MIC=100-625 ng/mL, MBC=312.5-1250 ng/mL) and indicated that the double bond at C-24 and C-25 positions in WU-FA-00 structure has little effect on its antibacterial activity. On the other hand, both WU-FA-00 and WU-FA-01 were more effective against Gram-positive strains of *Staphylococcus* than the Gram-negative strains, and this result is in accordance with the previous agar disk diffusion studies and implied that WU-FA-01 could be developed as an active antibacterial agent.

**Table 2.**
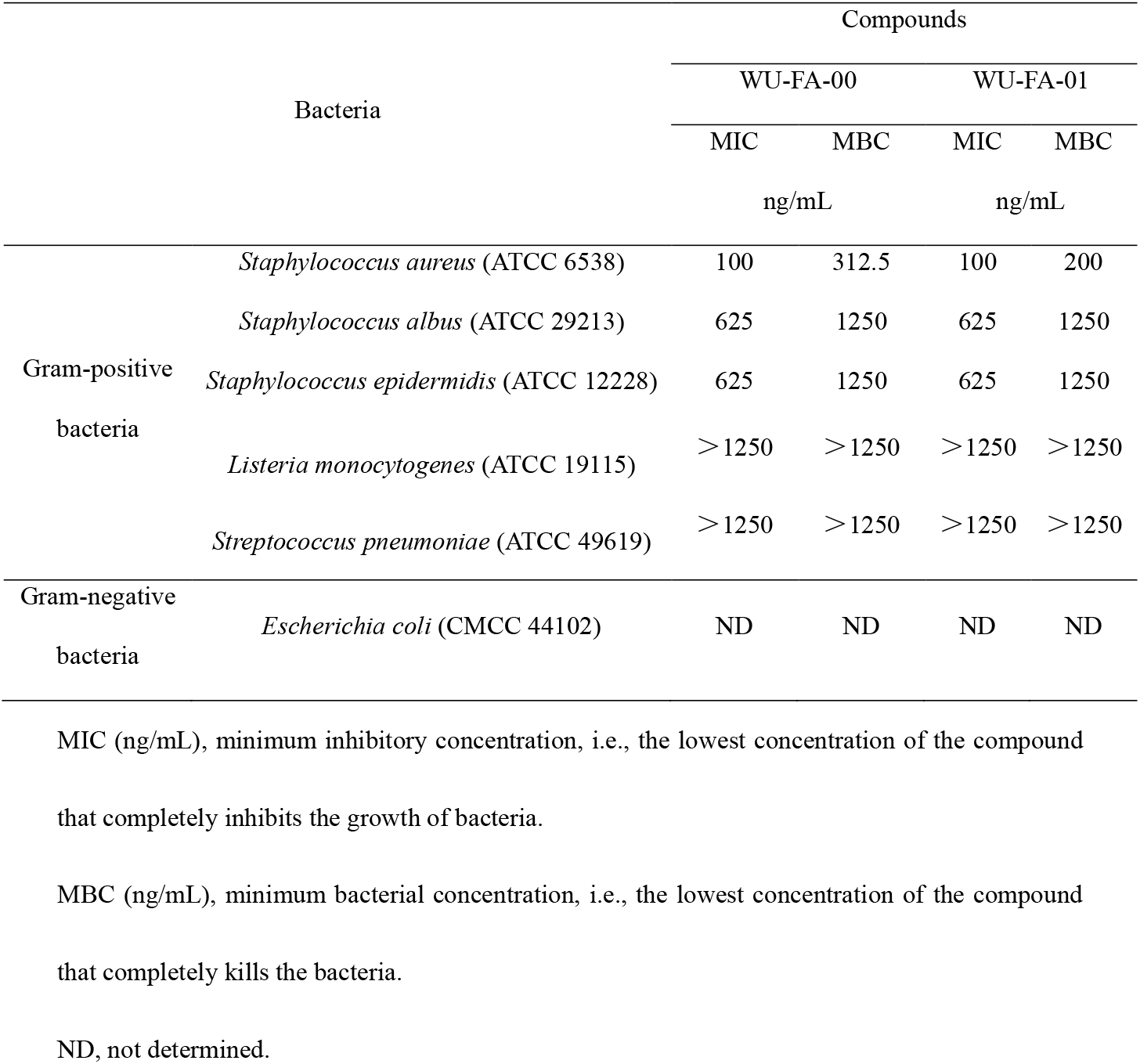
Antibacterial activities of WU-FA-00 and WU-FA-01 expressed in MIC and MBC (ng/mL).

### Killing kinetic studies

The time killing studies were carried out over a period of 24 h; bacteria were exposed to the tested compound at four different concentrations, which were determined according to their MICs. Figure 2 displays the time-kill curves of the tested compounds for *Staphylococcus aureus* (ATCC 6538), *Staphylococcus albus* (ATCC 29213), and *Staphylococcus epidermidis* (ATCC 12228). As shown in Figure 2, the MICs of the tested compounds were sufficient to inhibit almost all of the bacterial growth but with a slight increase after 20 h during this assay. Similar to the MBC results, no remarkable difference in the bacterial counts were found after incubation for 24 h at MICs, and the results confirmed that the MBCs were highly effective for killing bacteria. Furthermore, the bacterial population incubated with DMSO or with the test compounds, which were lower than that of their MICs, indicated less inhibitory action upon all selected bacterial strains. Moreover, Figure 2 also indicates that there is no difference in terms of killing kinetic between the two compounds against all of the chosen Gram-positive microorganisms.

**Fig. 2.**
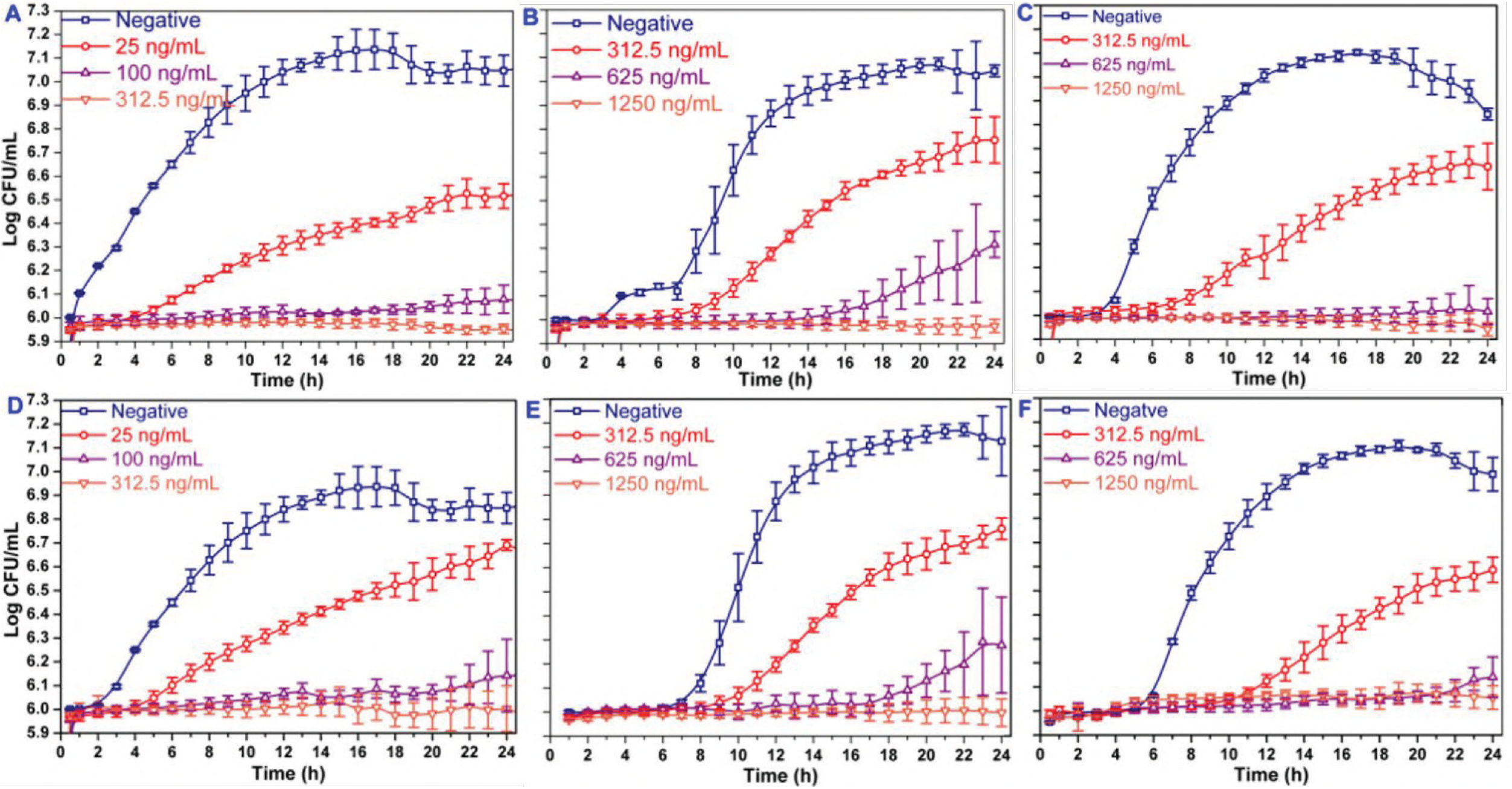
Time-kill curves for the three Gram-positive strains. Including *Staphylococcus aureus* (ATCC 6538) (A and D), *Staphylococcus epidermidis* (ATCC 12228) (B and E), and *Staphylococcus albus* (ATCC 29213) (C and F), exposed to four different concentrations of WU-FA-00 (Figure 2A, 2B, 2C) and WU-FA-01 (Figure 2D, 2E, 2F) according to their respective MICs (n=4).

### Inhibitory effects of WU-FA-00 and WU-FA-01 on TPA-induced edema in a mouse ear model

A TPA-induced ear edema mouse model was utilized to evaluate the *in vivo* anti-inflammatory activities of WU-FA-00 and WU-FA-01. It has been reported that TPA, which was normally adopted in this investigation model, is a well-known promoter of skin inflammation. The average weight of the ear punches is an important indicator that reflects the degree of skin edema when compared with the vehicle control group. As shown in Figure 3, the weight of mouse ear punches were significantly increased to 160.90% after 6 h when 20 μL TPA (0.125 μg/μL in acetone) was topically applied compared to the acetone-treated control group. Topical application of 2, 4 and 8 μg/mL of WU-FA-00 after TPA treatment modestly inhibited TPA-induced ear edema by 39.04%, 73.46%, and 83.83%, respectively, compared with the TPA group. However, 2, 4 and 8 μg/μL of WU-FA-01 significantly decreased the TPA-induced ear edema by 48.16%, 113.97% and 137.32%, respectively, in a dose-dependent manner. Furthermore, the compound WU-FA-01 had a similar effect on the positive control when it was used at a dose of 4 μg/mL (7.71 μmol/mL) with an inhibition rate of 113.97%, whereas dexamethasone had an inhibition rate of 134.13% at a dose of 2.5 μg/mL (6.37 μmol/mL). This result also suggested that WU-FA-01 had stronger protective effects than WU-FA-00 against TPA-induced skin inflammation.

**Fig. 3.**
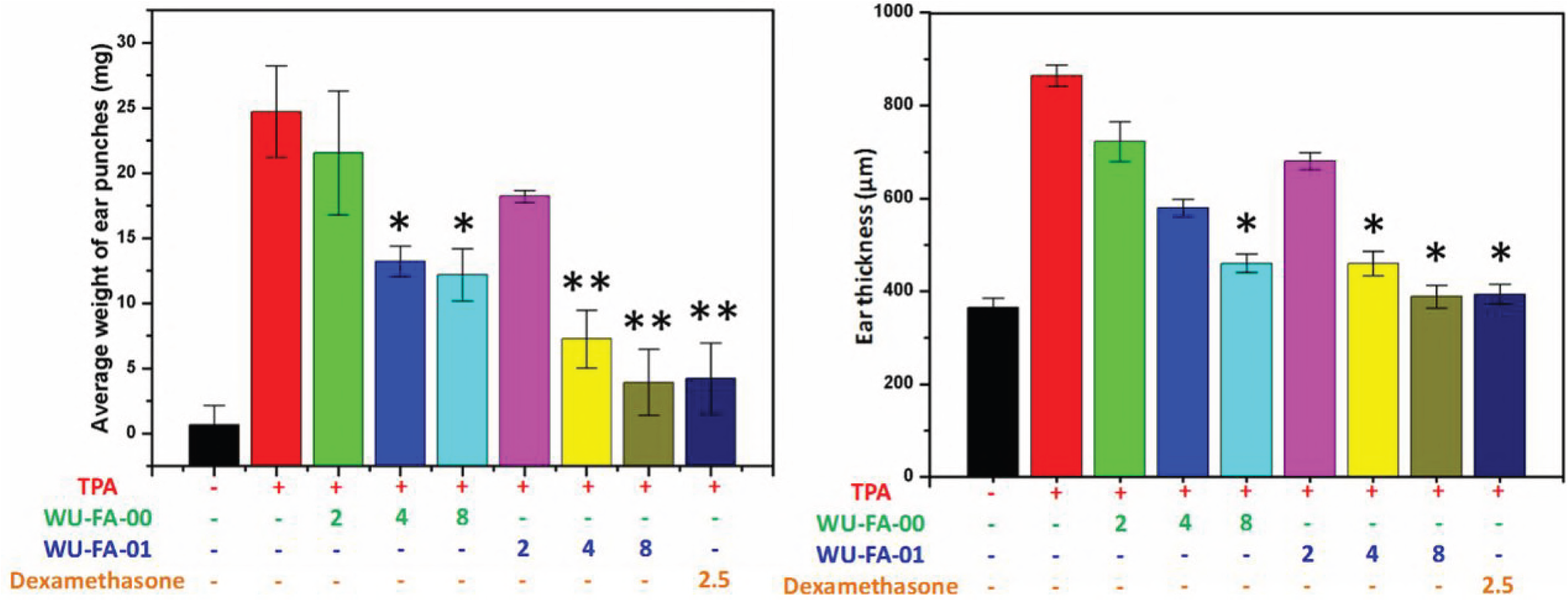
Inhibition effects of WU-FA-00 and WU-FA-01 on TPA-induced edema in mouse ears. The right ears of all animals (n=6) were topically treated with 20 μL of acetone (vehicle control) or WU-FA-00/WU-FA-01 (2, 4 or 8 μg/μL) in 20 μL of acetone after the application of TPA (2 μg/μL) in 20 μL of acetone. The mice were sacrificed 6 h after the TPA treatment. Both ear punches (9 mm in diameter) were immediately taken, and then, they were weighted and measured. The data from each group are expressed as the mean ± S.D. Compared to the TPA induced model group (n=6), \**P*<0.05, \*\**P*<0.01 (Dunnett’s multiple comparison test).

### Inhibitory effects of WU-FA-00 and WU-FA-01 on the histological appearance of mouse ears

To investigate the role of WU-FA-00 and WU-FA-01 plays in the histological appearance of a TPA-induced mouse ear model, both WU-FA-00 and WU-FA-01 were evaluated by transdermal application. In this model, the right ears of each group of mice were pretreated with 20 μL TPA (0.125 μg/mL in acetone), while the controls were topically adopted with acetone. The treatment compounds (20 μL) at three different concentrations were dissolved in acetone and used 5 min later. Dexamethasone was used as a positive control at a concentration of 2.5 μg/mL (6.37 μmol/mL) in acetone. After the ear tissues had been stained with H&E stain, as shown in Figure 4, the histological appearances of the ear sections indicated that the ears treated with acetone alone appeared normal in the epidermal layer without any obvious lesion. However, the TPA alone group displayed significant swelling, which was consistent with the results of the ear thickness and the ear punch weight (Figure 3). Moreover, the topical application of WU-FA-00 and WU-FA-01 could effectively suppress signs of the inflammatory response, such as epidermal hyperplasia and dense dermal leukocyte infiltration.

**Fig. 4.**
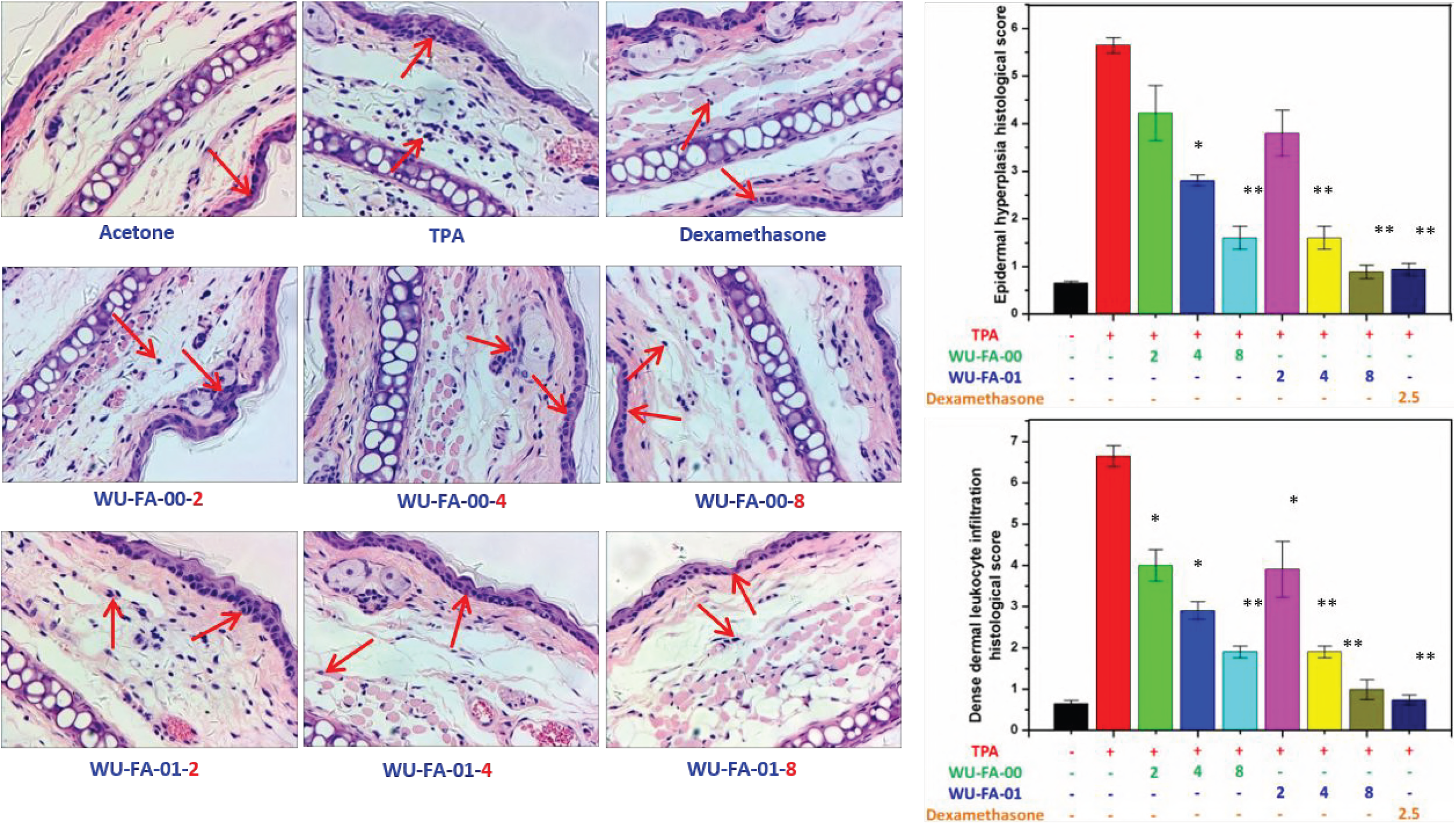
H&E staining for histological changes of TPA-induced mouse ears treated with an acetone control, TPA, WU-FA-00 and WU-FA-01 at different concentrations. The data from each group are expressed as the mean ± S.D. Compared to the TPA-induced model group (n=6), \**P*<0.05, \*\**P*<0.01 (Dunnett’s multiple comparison test). Magnification 200×.

### Inhibition of TPA-induced expression of TNF-α, IL-1β, COX-2

To gain insight into the molecular mechanisms by which WU-FA-00 and WU-FA-01 suppressed TPA-induced skin inflammation, we examined the effects of both WU-FA-00 and WU-FA-01 on the expression levels of pro-inflammation cytokines, including TNF-α, IL-1β and COX-2, in mouse ears using immunohistochemical analysis. As shown in Figure 5, the expression level of pro-inflammation cytokines (TNF-α, IL-1β and COX-2) were dramatically elevated 6 h after topical stimulation with TPA, which was apparently down-regulated in a dose-dependent manner by treatment with WU-FA-00 and WU-FA-01. However, the pro-inflammation cytokines levels of TNF-α, IL-1β and COX-2 between the treated groups and control group in the TPA-induced mouse ear model, were increased 20.37, 31.47 and 3.16-fold. Firstly, 2, 4 and 8 mg/mL of WU-FA-00 retarded TPA-induced overexpression of TNF-α by 5.1%, 52.9% and 80.7%, while 2, 4 and 8 mg/mL of WU-FA-01 retarded TPA-induced overexpression of TNF-α by 20.5%, 56.5% and 82.5% relative to the TPA group, respectively (Figure 5A). Secondly, WU-FA-00 at 2, 4 and 8 mg/mL greatly reduced the overexpression of IL-1β by 36.0%, 59.8% and 86.1%, while 2, 4 and 8 mg/mL of WU-FA-01 greatly reduced the overexpression of IL-1β by 35.9%, 65.6% and 86.6%, respectively (Figure 5B). Thirdly, 2, 4 and 8 mg/mL of WU-FA-00 retarded TPA-induced overexpression of COX-2 by 8.3%, 26.7% and 45.8%, while 2, 4 and 8 mg/mL of WU-FA-01 retarded TPA-induced overexpression of COX-2 by 12.1%, 31.6% and 56.6%, compared to the TPA group, respectively (Figure 5C). Therefore, the above results indicate that WU-FA-00 and WU-FA-01 could markedly suppressed the overexpression of pro-inflammation cytokines, which was in accordance with the previous results of ear weight and ear thickness (Figure 3) and histological changes (Figure 4) in this TPA-induced ear model.

**Fig. 5.**
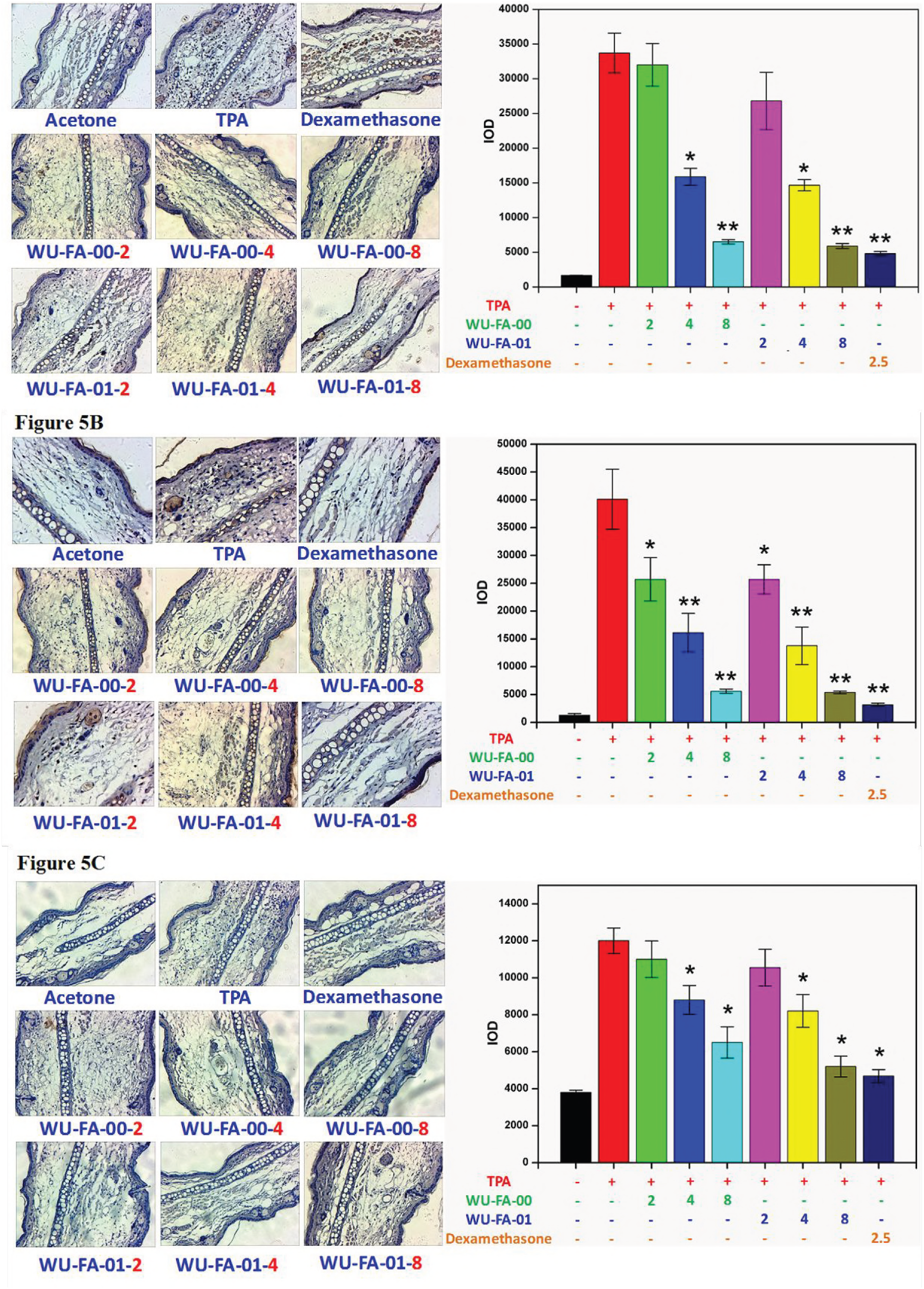
Effects of WU-FA-00 and WU-FA-01 treatment on pro-inflammation cytokines levels of TNF-α (Figure 5A), IL-1β (Figure 5B) and COX-2 (Figure 5C) in a mouse ear model. Mouse ears treated with acetone, TPA, WU-FA-00 and WU-FA-01 at different concentrations were analysed by immunohistochemical staining. The data are shown as the mean±SD. Compared to the TPA-induced model group (n=6), \**P* < 0.05, \*\**P* < 0.01 (Dunnett’s multiple comparison test). Magnification 200×.

### Inhibition of TPA-induced expression of p65, IκB-α, and p-IκB-α

The activation of NF-κB is significant for the regulation of TNF-α, IL-1β and COX-2 overexpression in the TPA-induced inflammatory model. Therefore, whether WU-FA-00 and WU-FA-01 could affect the NF-κB signalling pathway was determined by immunohistochemical analysis. As illustrated in Figure 6A, the results revealed that p65 was markedly suppressed by the treatment of WU-FA-00 and WU-FA-01, in which both WU-FA-00 and WU-FA-01 were more active at a concentration of 8 mg/mL. Moreover, the results also confirmed that the transcriptional activity was markedly up-regulated in the TPA-induced model but was inhibited by WU-FA-00 and WU-FA-01 at 8 mg/mL.

**Fig. 6.**
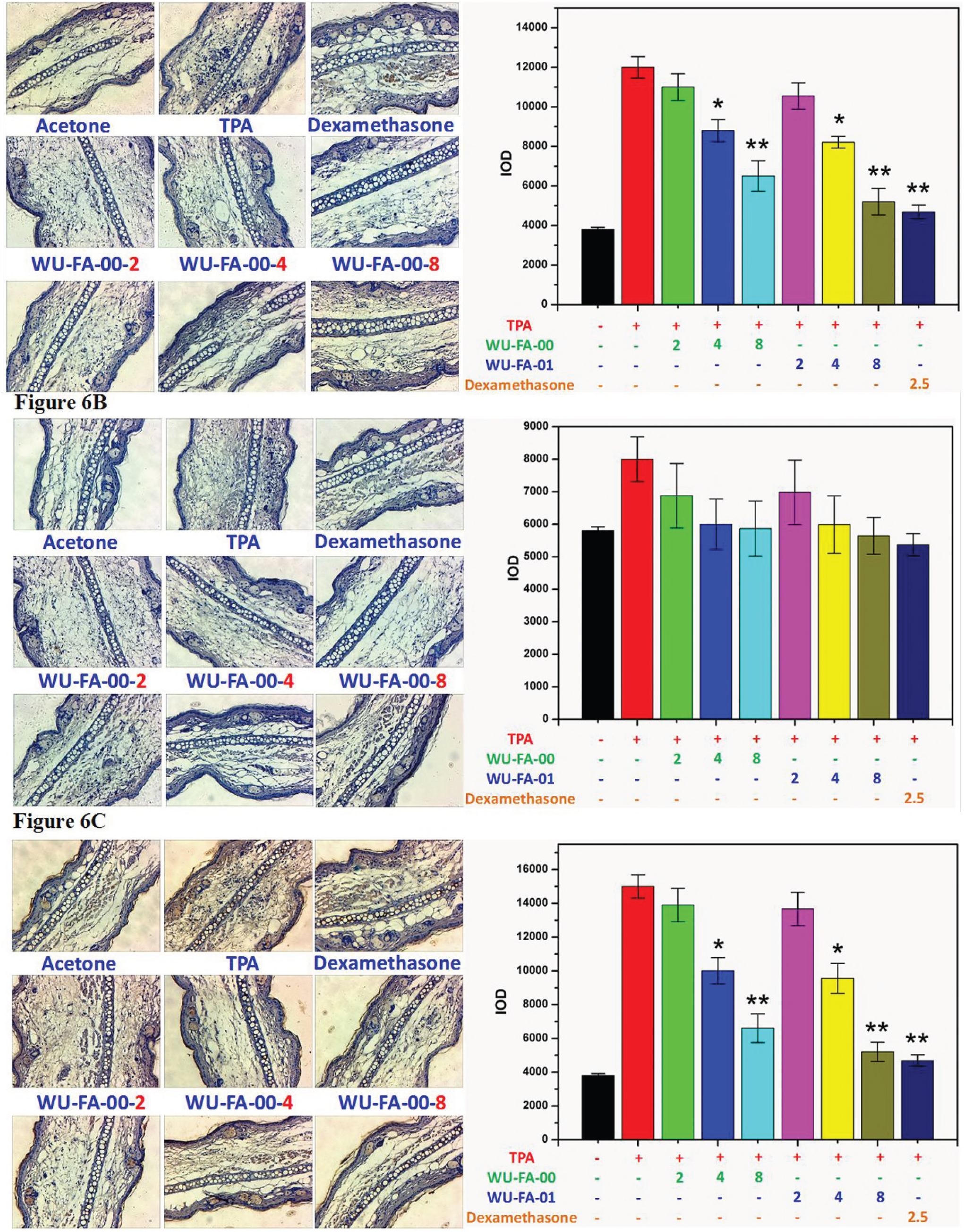
Inhibitory effects of WU-FA-00 and WU-FA-01 on the phosphorylation of p65 (Figure 6A), IκB-α (Figure 6B) and p-IκB-α (Figure 6C) in the TPA-induced ear model. Mouse ears treated with acetone, TPA, WU-FA-00 and WU-FA-01 at different concentrations were analysed by immunohistochemical staining. The data are shown as the mean±SD. Compared to the TPA-induced model group (n=6), \**P*< 0.05, \*\**P*<0.01 (Dunnett’s multiple comparison test). Magnification 200×.

The signalling pathway of IKK is involved in the induction of pro-inflammation cytokines *via* the modulation of NF-κB. Thus, it is necessary to gain insights into the IκB-α/p-IκB-α pathway in this TPA-induced model. From the immunohistochemical analysis in Figure 6B and Figure 6C, the levels of IκB-α and p-IκB-α in the TPA group were significantly increased. However, they could be suppressed by WU-FA-00 and WU-FA-01 in a dose-dependent manner, especially at a higher concentration of 8 mg/mL. These results imply that both WU-FA-00 and WU-FA-01 might block the activation of NF-κB through interfering with p65 and IκB-α/p-IκB-α to inhibit the expression of the TPA-induced pro-inflammation cytokines of TNF-α, IL-1β and COX-2.

## Discussion

Both WU-FA-00 and WU-FA-01 not only possessed excellent *in vitro* antimicrobial activities for Gram-positive *Staphylococcus* strains but also exhibited effective inhibition effects in the TPA-induced mouse ear model. Thus, both WU-FA-00 and WU-FA-01 could be considered as inhibitors of inflammation induced by bacterial infection. Moreover, it is possible that inflammation is frequently triggered by bacterial infection. The inhibitory effect of WU-FA-01 against microorganisms and TPA-induced skin inflammation is similar to its parent compound. The inhibitory effect of both compounds was associated with the suppression of TPA-stimulated pro-inflammation cytokines of TNF-α, IL-1β and COX-2. This study provides a further understanding of the anti-inflammatory properties of WU-FA-00 and WU-FA-01. Therefore, the results of this study implied that fusidic acid and the dihydro-analouge could be developed as di-functional agents, which possess both antimicrobial and anti-inflammatory activities.

## Methods and materials

### Chemicals

Fusidic acid (FA) was purchased from Macklin Co., Ltd., with over 98% purity. 12-0-tetradecanoylphorbol-13-acetate (TPA) was ordered from Sigma-Aldrich Chemical Co. (Saint Louis MO). TPA, FA and its derivative were dissolved in acetone to produce the desired concentrations of each compound. TNF-α and IL-1β antibodies were purchased from Bioss biotechnology Co. (Beijing, China) and Beyotime Biotechnology Co. (Beijing China). The silica gel (200-300 mesh) used in the column chromatography was supplied by Inno-chem Co., Ltd. (Beijing China). All other reagents and solvents were purchased from Adamas Reagent Ltd. (Shanghai China) or other commercial suppliers in their analytically or chemically pure forms and used without purification. TLC was performed on pre-coated silica gel F_254_ plates (0.25 mm; E. Merck); the starting material and the product were detected by either viewing under UV light or treating with an ethanolic solution of p-anisaldehyde spray followed by heating. The antimicrobial activity was determined by using a Multi-model Plate Reader (Infinite 200).

### Preparation of WU-FA-01

A 100-mL glassware was flamed-dried and allowed to cool in a desiccator before use. FA (1.0 g, 1.94 mmol) was dissolved in 50 mL of ethanol. 5% palladium on calcium carbonate (0.1 g, 0.19 mmol) was added to the reaction. Moreover, the reaction mixture was subjected to a vacuum-nitrogen purge and left to stir under a hydrogen atmosphere for 3 h. TLC was eluted in the mixture of Ethyl acetate: Petroleum ether=1:2 (V:V) and stained in *p*-anisaldehyde. Rf values of the starting material were 0.14, and the product was 0.17. Then, the reaction mixture was filtered through a pad of Celite and washed with ethyl acetate. The solvent was removed under vacuum to obtain a white solid. Yield: >98%.

### Microorganisms and culture conditions

Six bacterial strains were used for the bioassays, including three Gram-positive species, *Staphylococcus aureus* (ATCC 6538), *Staphylococcus albus* (ATCC 29213), *Staphylococcus epidermidis* (ATCC 12228), *Listeria monocytogenes* (ATCC 19115), and *Streptococcus pneumoniae* (ATCC 49619), and one Gram-negative species, *Escherichia coli* (CMCC 44102). All bacteria were maintained on Mueller-Hinton agar, and the cultures were stored at 4 °C and sub-cultured every week.

### Agar disk diffusion method

The antimicrobial activity of WU-FA-00 and WU-FA-01 were determined according to the standard agar disk diffusion method with a slight modification (24–26). A 0.5 McFarland (1×10^7^ to 1×10^8^ CFU/mL) concentration of the bacterial suspension was uniformly inoculated onto Mueller Hinton agar (MHA) solidified in 120 mm Petri dishes. Once the dishes were prepared, 6 mm-diameter discs of filter paper containing 5 μL of the examined compound, which had been diluted ten times with dimethyl sulfoxide (DMSO), were pressed gently against the surface of the agar. Discs containing WU-FA-00 were used as the positive control, while DMSO was used as the negative control. The dishes were incubated in a constant temperature incubator at 37 °C for 24 h. The inhibition zone (IZ) diameter was measured by a vernier caliper. All of the experiments were performed in triplicate.

### Broth microdilution method

The minimum inhibitory concentration (MIC) and the minimum bactericidal concentration (MBC) were determined by a microdilution method in 96-microwell plates according to Clinical and Laboratory Standards Institute (CLSI), with a slight modification (27, 28). A dilution series of the test compounds were obtained with DMSO as the solvent by two-fold serial dilution. The final concentrations of the test compound were 1~400 μg/mL. Each well received 5 μL of a specific concentration of the compound and 195 μL of Mueller Hinton broth inoculated with the test microorganism (1.5×10^5^ CFU/mL); the final concentration of the test compound reached 0.025~10 μg/mL. WU-FA-00 and DMSO were treated as a positive control and a negative control, respectively. The microplates were incubated in a bacteriological oven for 24 h at 37 °C, and the drug susceptibility results were monitored by measuring the absorbance at 600 nm using a Multimodel Plate Reader (Infinite 200). The lowest concentration without visible growth was defined as the MIC.

The minimum bactericidal concentrations (MBCs) were determined based on the MIC results (29, 30): serial sub-cultivation of a 5 μL aliquot near the MIC in microtiter plates containing 195 μL of Mueller Hinton broth per well; incubation for 24 h at 37 °C. The lowest concentration of antimicrobial agent that killed at least 99.9% of the starting inoculum was defined as the MBC endpoint, which was determined as the lowest concentration with no visible growth by measuring the absorbance at 600 nm using a Multimodel Plate Reader (Infinite 200). All experiments were conducted in triplicate.

### Killing kinetic studies

The killing kinetic assay on the Gram-positive strains (27, 31, 32), including *Staphylococcus aureus* (ATCC 6538), *Staphylococcus albus* (ATCC 29213), and *Staphylococcus epidermidis* (ATCC 12228), was performed against WU-FA-00 and WU-FA-01 in 96-microwell plates, respectively, and four different concentrations (0, 25, 100, 312.5 ng/mL) of each compound were tested. The microplates were incubated for 24 h at 37 °C, and the growth of bacteria was monitored by measuring the absorbance at 600 nm using a Multimodel Plate Reader (Infinite 200) every 1 h.

### Animals, diets and treatments

Female Kunming mice approximately 22-25 g were used in the TPA-induced *in vivo* model. All animals were supplied by the Experimental Animal Centre of Guangdong Province. They were maintained at 25±1°C with standard mouse chow diet and tap water *ad libitum* and were kept on a regular light-dark cycle with 50% relative humidity. All the animal experiments were performed according to the Ethical Regulations on Animal Research of Southern Medical University (Approval Documents: SCXK/20130002).

### TPA-induced skin inflammation in mouse

The mice were divided into nine groups: each group consisted of six mice, including a blank group, a TPA group, a dexamethasone group, and six groups for WU-FA-00 and WU-FA-01. In the mouse ear edema model, 20 μL of acetone vehicle was topically applied to the right ear, and 20 μL of the treatment compounds at three different concentrations, which were dissolved in acetone, were used 5 min later after 20 μL of TPA (0.125 μg/mL in acetone) was previously applied to induce the inflammation model (33, 34). Dexamethasone at a concentration of 2.5 μg/mL (6.37 μmol/mL) in acetone was used as the positive control. Then, all of the mice were maintained at a standard condition and sacrificed 6 h after TPA treatment. Two ear punches (9 mm in diameter) from the right and left ears were then harvested immediately and weighted; the left ear was used for comparison. All experiments were carried out in compliance with the relevant laws and institutional guidelines, which were all approved by the Southern Medical University (Approval Documents: SCXK/20130002).

### Histological appearance of mouse ears

The right ear punches were fixed in 10% neutral buffered formalin, decalcified in EDTA buffer, subjected to a series progression of dehydration and embedded in paraffin. Sections of 9 mm were cut by using a microtome and were mounted on colourfrost microslides (VWR scientific, Edmonton, Alberta, Canada). The sections were dried overnight and stained with haematoxylin and eosin (H&E) in accordance with the classical methods of histology. Images of the sections representing each treatment group were observed under a microscope (Olympus, Japan) to evaluate the damage of ear tissue.

### Scoring the expression of biomarkers

Each histologic type of lesion in the TPA-induced ear model was scored independently by two experienced investigators who were not aware of the identity of the specimens (×200) (33, 35). The staining intensity was scored as follows: 0, no staining; 1+, faint; 2+, moderate; and 3+, strong.

1+, 2+ and 3+ were recorded as 1, 2 and 3 points, respectively. The staining extent was graded as follows: 0, no staining; 1+, ⩽25% of cell positive; 2+, 26% to 50% of cells positive; and 3+, ⩾51% of cells positive.

### Immunohistochemical detection of TNF-α, IL-1β, COX-2, p65, IκB-α, and p-IκB-α expression

The ear punch tissues were fixed in formalin, and paraffinised sections of 5 μm thickness were incubated with 1.2% H_2_O_2_ in PBS to quench the endogenous peroxidase activity in order to minimize nonspecific staining. Then, the sections were washed three times (5 min each) with 1X TBST (0.05% Tween-20). Subsequently, the primary antibody of a proliferating cell nuclear antigen was diluted 100 times, applied to each section and left overnight at 4 °C. The sections were washed with PBS and incubated with a biotin-conjugated horseradish peroxidase antibody (1:200) for 1 h at room temperature. Finally, peroxidase was detected using the 3, 3-diaminobenzidine tetrahydrochloride reaction, which produced a brown label in the epidermal tissue. The cells that stained positive for TNF-α, IL-1β and COX-2 were counted in the section of the mouse ear using the Image-Pro Plus (Version 6.0) software (33). The results were expressed as the number of stained cells. Immunohistochemical analysis of p65, IκB-α and p-IκB-α were also conducted to gain insight into the signalling pathway of WU-FA-00 and WU-FA-01 in the TPA-induced mouse ear edema model.

### Statistical analysis

The results are expressed as the mean ± standard error (SE) or standard deviation (SD). Statistical comparisons among groups were performed by using Dunnett’s multiple test. Statistical significance was defined by a *P* value of < 0.05.

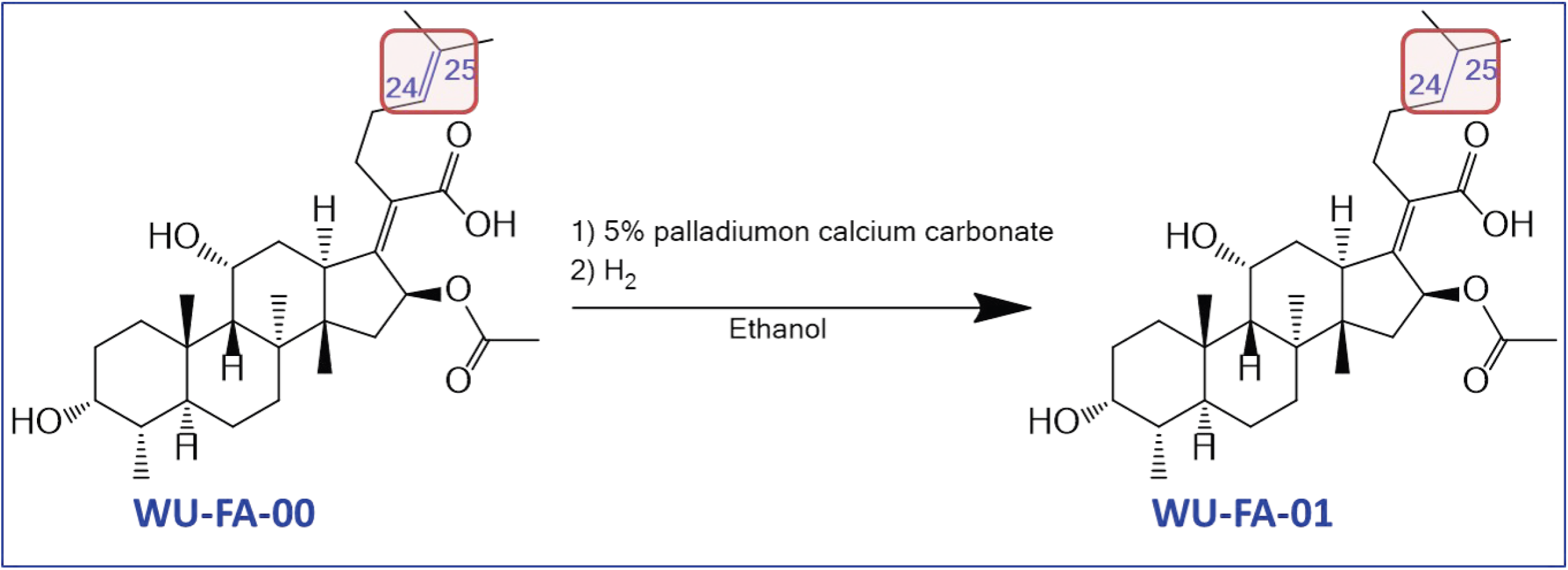

## Author Contributions

Performing the experimental work and drafting the manuscript: (PPW, XPC). Performing the bioactivity test: (PPW, HH, TRW, BRT, YYZ). Performing the experimental statistical analysis (PPW, XPC, MG, DLL, JZ, ZJS). The director as well as the designer of the manuscript: (WDH, XTX, KZ). The project coordinator: (WDH, SAW, PMO, SQZ).

## Acknowledgments

Sincere and heartfelt thanks must go to Miss Sulian Liang who has given generous suggestions on the English language of the manuscript.

**Scheme 1 Synthesis of the hydrogenation derivative of fusidic acid (WU-FA-01)**.

